# Disentangling polymer confinement from specific-folding interactions reveals the drivers of *E. coli* chromosome organization

**DOI:** 10.64898/2026.05.20.726692

**Authors:** Pourya Delafrouz, Tanya Nesterova, Frances Harris, Hammad Farooq, Lin Du, Jian Liu, Ao Ma, Jie Xiao, Jie Liang

## Abstract

The three-dimensional organization of the bacterial chromosome is critical for gene regulation. Chromosome conformation capture (Hi-C) has enabled genome-wide mapping of chromosomal folding, yet ensemble-averaged contact maps entangle biologically specific-folding interactions (SFIs) with nonspecific polymer compaction and mask single-cell heterogeneity. Here, we developed a polymer-based simulation framework in *E. coli* to address these limitations. We found that a null model of 300,000 random polymer configurations recapitulated the global chromosomal organization observed in Hi-C data, establishing that most Hi-C signals reflect generic polymer behavior in a confined volume. Contrasting null-model predictions with experimental Hi-C data isolated a small subset of SFIs (< 7%) and generated a specific-fold ensemble of 20,000 single-cell conformations that reproduced chromosomal interaction domains and single-cell heterogeneity. SFIs were enriched in the *ter* region, reduced nucleoid accessibility, colocalized with cryptic prophages, and depleted in positively supercoiled regions. H-NS and MatP emerged as major chromosome-wide and local determinants of SFIs, respectively. Furthermore, high-SFI regions correlated with stress-adaptive genes, whereas low-SFI regions harbored housekeeping genes. Together, our results established that a small number of biologically encoded SFIs superimposed on a polymer background shape the *E. coli* chromosome and gene expression, providing a quantitative framework for dissecting chromosome architecture and function.

## Introduction

The three-dimensional (3D) organization of bacterial chromosomes is critical for understanding how cells regulate gene expression, coordinate DNA replication, and respond to environmental stress (1). In *E. coli*, the chromosome consists of a single circular DNA molecule compacted into a protein-DNA-RNA macromolecular complex termed the nucleoid. The compaction results from both passive confinement within the cell membrane and active processes involving DNA supercoiling (2), replication and transcription (1), and interactions with structural maintenance of chromosome proteins (SMCs, (3)), nucleoid-associated proteins (NAPs, (4)), topoisomerases (5), and other DNA binding factors (6), forming a highly organized yet dynamic structure (2).

Experimental approaches, especially chromosome conformation capture techniques like Hi-C, have revealed fundamental organizational principles of the *E. coli* chromosome, including the formation of chromosomal interaction domains (CIDs), large-scale compartmentalization into left and right arms, macrodomains, and specific long-range interactions between regulatory loci (4, 7–9). These studies support the prevailing view that bacterial chromosome organization arises from a combination of entropy-driven polymer dynamics and enthalpy-driven molecular interactions mediated by NAPs such as HU, MatP, and H-NS (10–12). However, robust single-cell Hi-C remains essentially inaccessible in bacteria due to the minuscule amount of chromosomal DNA in each cell. Bacterial Hi-C measurements represent ensemble averages over millions of cells and obscure the intrinsic heterogeneity of individual chromosome conformations in single cells, which may give rise to phenotypic and functional heterogeneities at the population level. Advanced superresolution imaging approaches can resolve global chromosomal conformations in individual cells but lack the throughput and coverage required to reconstruct full chromosome-level architectures at the single-cell level (13).

Computational modeling offers a powerful framework for simulating chromosome configurations from first principles of physics, enabling the integration of single-cell imaging data with ensemble-averaged sequencing measurements from millions of cells. Polymer-based models have demonstrated that much of the large-scale organization observed in Hi-C maps can be captured by generic polymer behavior and excluded-volume effects (14–16). However, these models often relied on parameter fitting or the direct imposition of Hi-C constraints, making it difficult to discern which structural features emerge intrinsically from confinement and polymer dynamics, and which arise from biologically mediated specific-folding interactions (SFIs). Furthermore, extensive modeling efforts have focused on linear eukaryotic chromosomes (17–19), leaving the distinct topology and spatial constraints of circular bacterial chromosomes underexplored.

In this work, we addressed these limitations by developing a polymer-based chain-growth simulation framework tailored to the *E. coli* chromosome and readily generalized to circular chromosomes of other bacterial species. This framework explicitly accounts for the genome’s circular topology and confinement within the nucleoid, thereby generating large ensembles of single-cell chromosome conformations driven exclusively by entropy and excluded-volume polymer dynamics. The resulting structure, devoid of any specific enthalpic molecular interactions, defines a “null model” for chromosome organization. By comparing the chromosomal contact map derived from this null model to experimental Hi-C data, we can distinguish polymer-driven, nonspecific contacts from biologically mediated SFIs, thereby disentangling the entropic and enthalpic contributions to average chromosome organization. Moreover, by statistically decomposing the ensemble into distinct, mutually exclusive single-cell chromosome conformations, we can recover the intrinsic heterogeneity of specific chromosome folding at the single-cell level, which is experimentally inaccessible in bacteria. We have previously successfully applied this strategy to identify specific chromosome folds in *Drosophila* (18) and human cells (17, 20, 21), demonstrating our framework’s general applicability across organisms.

Using this strategy, we demonstrated that excluding entropic, nonspecific chromosomal interactions was essential for isolating SFIs, and that SFIs are the drivers of *E. coli* chromosome organization, as they define chromosomal interaction domains, establishing insulation boundaries, and revealing region-specific compaction patterns. Our analyses further highlighted how organizational heterogeneity shaped differential chromosome accessibility across the *ori* and *ter* regions and how highly ranked SFI regions were preferentially associated with cryptic prophages, specific nucleoid-binding proteins, and transcriptional repression. Together, these findings provide new mechanistic insights into how *E. coli* chromosome organization emerges from the interplay between stochastic polymer physics and biologically encoded molecular interactions, advancing our understanding of bacterial chromosome organization.

## Method

### Polymer model setup

The model represents the *E. coli* chromosome as a circular, coarse-grained polymer, with each bead corresponding to a 5-kb genomic segment. The chromosome was confined within a capsule-shaped volume that approximates the shape and dimensions of an *E. coli* nucleoid (2.65 µm by 0.9 µm). Simulations were implemented in C++ and optimized with OpenACC for parallel execution on NVIDIA GPUs, enabling efficient sampling of large structural ensembles. Chromosome conformations were generated via Monte Carlo sampling, yielding a diverse ensemble of single-cell structures that capture stochastic variability and the effects of specific folding interactions (SFIs). Our analysis shows that these specific interactions are caused by sequence-specific binding and remodeling activity that shape chromosome architecture. The simulation code, which will be made available on GitHub, is an enhanced version of a previously developed model for *Drosophila* (18), restructured to incorporate *E. coli*-specific features such as the circular DNA topology and capsule-shaped confinement, and further optimized for GPU-based parallel performance. When constructing Hi-C contact frequency maps from chromosome configurations, we defined the interaction probability as the fraction of conformers in which the surface-to-surface distance between a pair satisfies d < 80 nm. The generated conformations using SFIs preserve the native genomic coordinates, where each bead corresponds to a defined genomic interval (e.g., bead 1: 0–5 kb, bead 2: 5–10 kb). This enables direct mapping of genomic features such as the *ori* and *ter* regions onto the 3D chromosome conformations.

### Hi-C data processing

We aligned Hi-C sequencing reads to the *E. coli* reference genome (NC-000913.3) and processed them using the HiC-Pro pipeline (v3.1.0) (22). The dataset from Cockram et al. (8), with accession number PRJNA587586 (BioProject Number) and SRR10394904 (run number; HpaII digestion) was analyzed at 5-kb resolution. Contact count matrices were normalized using the iterative correction and eigenvector decomposition (ICED) method (23) and the normalized counts were subsequently log2-transformed after adding a pseudo-count.

### ChIP-seq and REP element data processing

We fetched data for all proteins mentioned in the analysis (**Figure 4**) from the NCBI database using SRA accession numbers provided in the main text. We ensured that all ChIP-seq data were collected during early exponential growth in minimal media so that it is comparable to the Hi-C map in Cockram et al. (8). Using Bowtie2 (24), we aligned the reads contained in the downloaded FASTQ files to the *E. coli* reference genome (NC-000913.3). Then, we created a merged and sorted BAM file using Samtools on the resulting SAM file from the previous step (25). Enriched regions (peaks) were identified in the BAM file using the callpeak function (26). In post-processing, we used a 5 kb bin size to determine protein log-enrichment score at each genomic position, which we then compared to percent coverage of each SFI per bin. To determine REP element bin occupancy across the reference genome (NC-000913.3), we utilized RNA-seq data obtained by a colleague in our lab and quantified sequence frequency within 5-kb bins (27).

### Chromosome geometry and nucleoid accessibility

We assessed chromosome accessibility from 3D genome conformations using alpha-shape–based polyhedral representations (28). For each single-cell conformer, we computed the Delaunay tetrahedralization of the convex hull of all polymer beads, and extracted the polyhedron defined by the alpha shape at a specified parameter (29). Beads near the polyhedral surface were designated as accessible regions. Accessibility frequencies were computed for both the random-fold null model and the specific-fold ensemble. To estimate accessibility for macromolecules of different sizes, we varied the alpha parameter and calculated the total accessible volume for spherical probes of diameter *D*_*probe*_ (nm).

### Positive and negative super coiling analysis

For the two ChIP-seq datasets (GapR-seq (30) for positive supercoiling and EcTopoI-seq (31) for negative supercoiling), raw sequence reads were first assessed for quality using FastQC, and adapter sequences were trimmed with Trimmomatic (32). The processed reads were then aligned to the *E. coli* reference genome (NC-000913.3) using Bowtie2 (24), and the resulting SAM files were converted to BAM format with Samtools (25). Enriched regions (peaks) were identified using the MACS2 callpeak function. Each 5 kb genomic bin in the Hi-C maps was classified as a positive supercoiling site (+SC) if it contained the center of at least one GapR peak, or as a negative supercoiling site (-SC) if it contained the center of at least one TopoI peak.

### RNA-seq analysis

Normalized RNA-seq data for *E. coli* (SRR2932637) (33) were obtained from the original study. Because read mapping, quantification, and normalization had already been performed against the *E. coli* reference genome (NC-000913.3), we used the processed expression matrix directly for downstream analysis.

### Perturbation of cryptic prophage

To test whether SFIs are enriched in cryptic prophage regions, we compared the overlap between 11 annotated cryptic prophage regions and 46 genomic bins classified as high-SFI regions. We then performed 10,000 random perturbations across the 928 genomic beads to estimate the expected overlap under random placement. The resulting null distribution was used to assess whether the observed overlap for high-SFI and low-SFI regions was greater than expected by chance.

## Results

### Modeling the *E. coli* chromosome as a circular, self-avoiding polymer

To investigate the conformation of the *Escherichia coli* K-12 chromosome (4,608 kb), we constructed a coarse-grained polymer model in which each polymer bead represented 5 kb of DNA and had an effective diameter of ∼ 67.5 nm, estimated from the reported DNA occupancy in the nucleoid volume of *Escherichia coli* (14). The entire chromosome was thus represented by 928 connected beads confined within a capsule-shaped nucleoid (2.65µm in length and 0.9 µm in diameter), based on our previous superresolution imaging measurements (34) (**Fig. 1A**).

**Figure 1:**
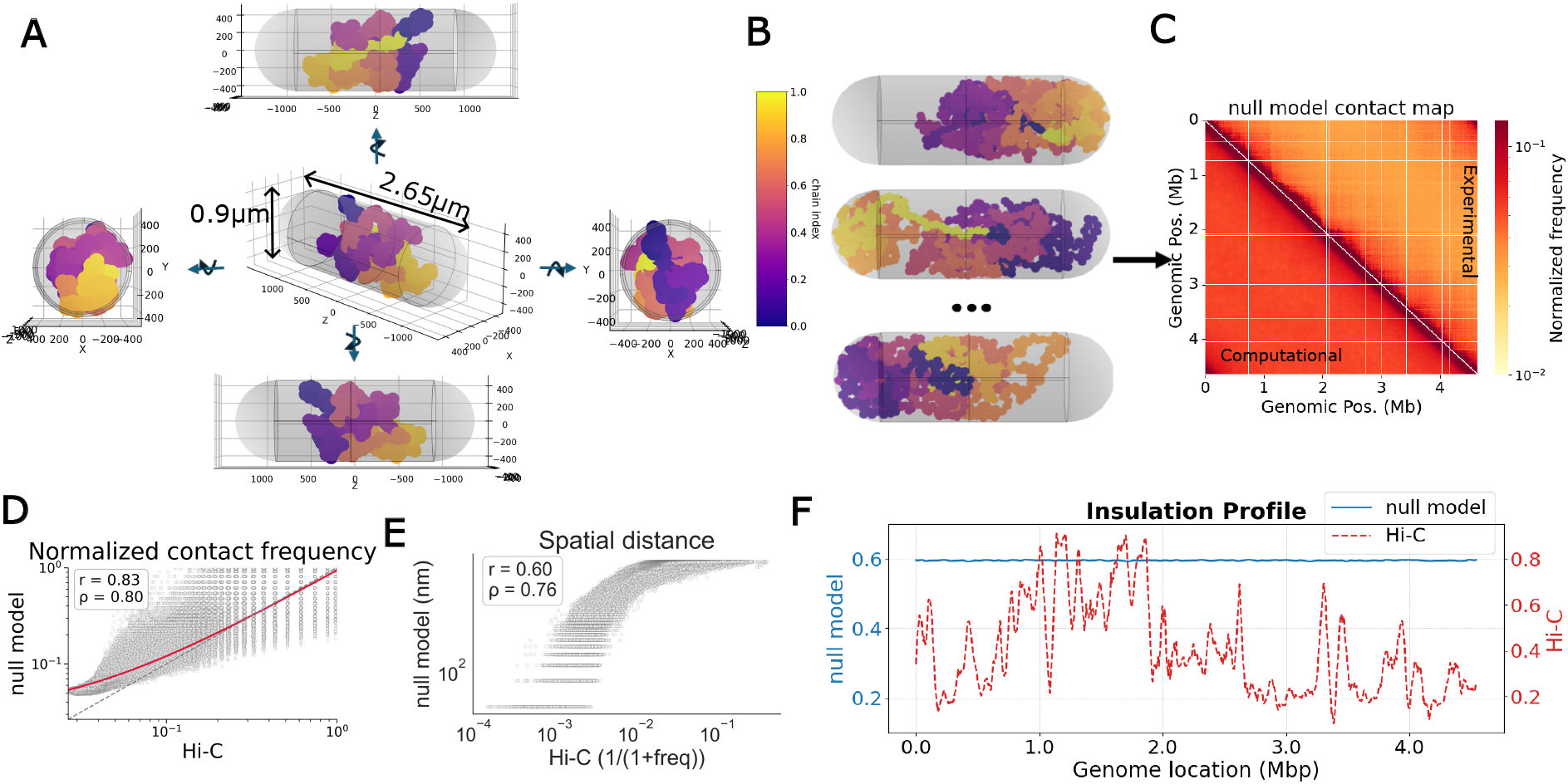
Simulated null model of *E. coli* chromosome conformation reproduces the overall chromosomal structure but not interaction domains. **A**) Dimensions of a capsule-shaped *E. coli* nucleoid (center, gray) and the simulated chromosomal polymer chain (colored from purple bead 1 to yellow bead 928, coarse-grained to 300 beads for clarity in visualization), with different views of the same configuration on the four sides. **B**) Examples of individual simulated random polymer chains in single cells (totaling 300,000 conformers). **C**) The contact map (bottom triangle) generated from the 300,000 random-fold conformers compared with the experimentally measured Hi-C map (top triangle). **D**) The contact frequency of the null model (y-axis) correlates significantly with the experimental Hi-C contact frequency (x-axis) based on pixel-wise comparison in **C. E**) The 3D spatial separation from the null model correlates significantly with the reciprocal of Hi-C contact frequency, which scales with the spatial separation. **F**) Insulation score analysis shows that the null model does not replicate chromatin interaction domain (CID) boundaries (blue) as observed in experimental Hi-C data (red).

The model treats the chromosome as a self-avoiding, random-walk chain of an appropriate length, with the first and last beads connected to ensure circular topology. The chain-growth process begins by randomly placing the first bead within the nucleoid volume, after which subsequent beads are added sequentially to available positions immediately adjacent to the preceding bead. A new bead is accepted if the current end-to-end distance (*d*) between the first and the new beads satisfies *d* < *N*_*r*_ × *D*_*b*_, where *N*_*r*_ is the number of remaining beads, and *D*_*b*_ is the bead diameter. At the completion of chain growth, the final bead position is accepted only if its distances to both the first and the preceding bead are equal to the bead diameter *D*_*b*_. Any chain that cannot be closed under these constraints is rejected. The random positioning of the first bead and the distance constraint during chain growth ensure that the resulting conformations preserve both the circular topology and stochasticity, thereby generating physically realistic chromosome conformations. Using this chain-growth process, we produced a chromosome-fold null model consisting of 300,000 random, independent *E. coli* chromosome conformers (**Fig. 1B**), capturing cell-to-cell variabilities observed in imaging studies of bacterial chromosome organization (35).

### Simulated chromosome-fold null model reproduces experimentally determined global organization

Using the simulated chromosome conformers, we evaluated whether the null model could replicate the overall structural features observed in an experimentally measured *E. coli* Hi-C map obtained from K-12 MG1655 cells grown at 37 °C in minimal medium (8). Here, we defined the interaction probability between any two beads in the simulated circular chromosome as the frequency with which their surface-to-surface distance was less than 80 nm across all conformers. We chose the 80-nm threshold based on crosslinking distance estimates from previous Hi-C studies (36, 37). We then compared the simulated, null model-based contact map and the experimental Hi-C contact map using ICE-normalized matrices in log space (**Fig. 1C**, Methods). A pixel-by-pixel comparison of the two contact maps showed a significant correlation (*r*_*Pearson*_= 0.83 and ρ_*Spearman*_ = 0.80, **Fig. 1D**). Furthermore, Hi-C derived pseudo separation distance (1/(1+contact frequency) also exhibited significant correlation with the 3D spatial separation calculated from the null model (*r*_*Pearson*_= 0.60 and ρ_*Spearman*_ = 0.76, **Fig. 1E**). These results suggested that the global chromosome fold is primarily driven by the chromosome’s inherent polymer behavior rather than by specific locus-locus interactions.

To further assess how accurately the simulated chromosome captured the overall folding pattern seen in experiments, we calculated the end-to-end distances of 2 Mbp segments at various genomic locations in the null model. The resulting distances remained nearly constant across different loci (**Fig. S1A**), reflecting the consistent trend observed in the corresponding Hi-C contact frequencies (**Fig. S1B**). Additionally, the spatial separations between all pairs of chromosome loci in the null model scaled as a power law of genomic separation (**Fig. S1C**), consistent with scaling relationships previously derived from experimental chromosomal Hi-C maps (30). These results demonstrate that the null model robustly recapitulates global chromosome organization, consistent with a uniform polymer-like folding behavior of the *E. coli* chromosome.

### The null model does not reproduce chromosomal interaction domain boundaries

Next, we tested the null model’s ability to capture the organization of chromosomal interaction domains (CIDs). We calculated the insulation score along the genome using a sliding-window method that identifies local minima in Hi-C interaction frequencies, thereby determining CID boundaries (38). We found that the insulation profile of the null model was mostly uniform across the chromosome (**Fig. 1F**, blue curve), suggesting a lack of prominent domain boundaries. In contrast, the experimental Hi-C data showed distinct minima corresponding to the expected macrodomains and CID boundaries (**Fig. 1F**, red curve). Therefore, although the null model captured the overall folding properties of *E. coli*’s circular chromosome, it did not reproduce the sharp insulation boundaries seen in the experimental data, indicating that specific, biologically encoded folding interactions are needed to form CIDs.

### Isolating specific-folding interactions (SFIs)

The ability of the null model to recapitulate the global chromosome folding pattern observed in experimental Hi-C data, yet its failure to identify chromosomal interaction domain (CID) boundaries, suggested that differences between the experimental and simulated contact maps could pinpoint specific, molecular factor-driven interactions responsible for unique chromosomal folds beyond those expected from random, polymer-driven dynamics.

To identify these interactions, we compared experimentally measured Hi-C contact frequencies with those generated from the null model. We identified chromosomal bin pairs with experimental contact frequencies significantly enriched relative to the null expectation (**Fig. 2A**, left) and classified them as specific-folding interactions (SFIs). These SFIs represent contacts that cannot be explained solely by polymer confinement and excluded-volume effects.

**Figure 2:**
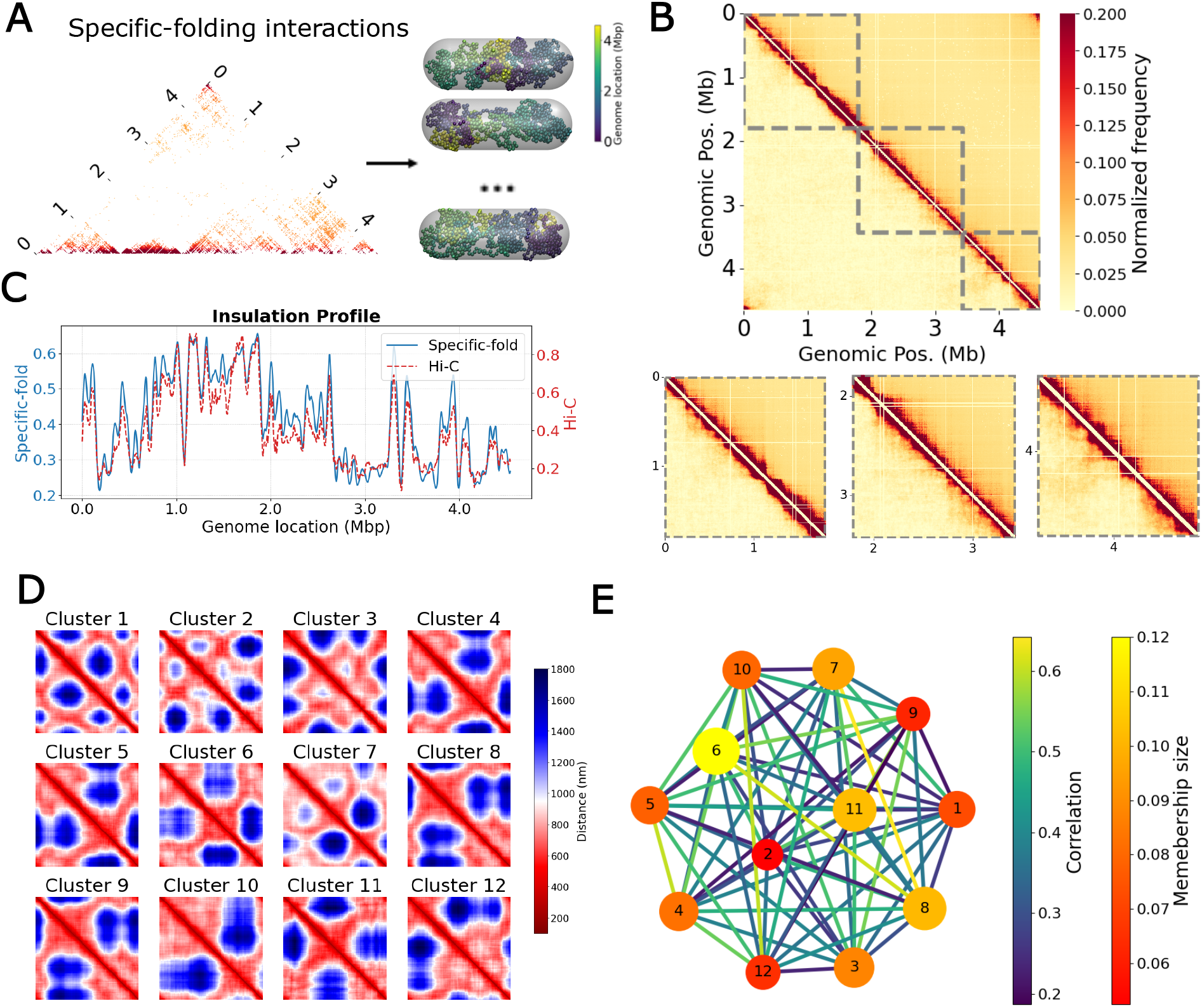
Individual *E. coli* chromosomes exhibit specific-folding interactions (SFIs) and heterogeneous conformations. **A**) Isolated specific-folding interactions (left) were used as spatial constraints to generate 20,000 single-cell conformations (right). **B**) Contact frequency map of 20,000 individual *E. coli* chromosome conformations simulated by the specific-fold model (bottom triangle) closely correlates (*r*_*Pearson*_ = 0.93, ρ_*Spearman*_ = 0.84) with the experimental Hi-C map (top triangle). Three zoomed-in heatmaps highlight individual CIDs (dashed grey boxes) that were captured by the specific-fold model. **C**) Insulation score for the specific-fold model (blue) and Hi-C experiment (red dashed line) are highly correlated (*r*_*Pearson*_ = 0.96). **D**) Average distance maps of twelve different clusters of single-chromosome conformations. **E**) Network map of each cluster (population percentage indicated by the size of the circle) and the Spearman’s correlation coefficients (colored lines) between each pair showed weak correlations.

Using SFIs as ensemble-level spatial constraints, we then applied a Markov Chain Monte Carlo (MCMC) approach to statistically sample a set of distinct, mutually exclusive single-cell chromosome conformations that collectively gave rise to the observed SFIs in the ensemble Hi-C map. These reconstructed single-cell chromosome conformations, which we termed the specific-fold model to contrast with the null model, contained biologically encoded structural features beyond those arising from random polymer folding, representing specific chromosome fold configurations in single bacterial cells that are currently inaccessible experimentally (**Fig. 2A**, right). We previously demonstrated the validity of this approach for identifying SFIs in eukaryotic chromosomes (17, 19, 20), providing a foundation for extending it to the circular *E. coli* genome.

Using the specific-fold model, we observed a strong pixel-by-pixel correlation between the simulated and experimental Hi-C contact maps (**Fig. 2B, Fig. S2A**, *r*_*Pearson*_= 0.93, *p*_*Spearman*_= 0.84). Furthermore, the insulation profile derived from the specific-fold model closely matched that of the experimental Hi-C contact map (**Fig. 2C**), thereby successfully reproducing the sharp CID boundaries of the *E. coli* chromosome.

Next, we investigated how these SFIs influenced the chromosome’s overall topology (39). Since these specifically folded single chromosomes have highly heterogeneous conformations (**Fig. S2B**), we applied k-means clustering to the distance maps of individual conformers and identified twelve distinct spatial clusters with varying population sizes (**Table S1**). Interestingly, the averaged distance map of each cluster revealed unique interaction patterns and differed significantly from other cluster maps (**Fig. 2D**), as demonstrated by the relatively moderate Spearman’s correlations between the average distance maps of these clusters (**Fig. 2E**). These single-chromosome conformation clusters are consistent with a previous imaging study, which mapped distances of chromosomal loci of single *E. coli* cells using seqFISH and identified nine distinct structural clusters (39). As shown in **Fig. S2C**, our single-cell conformation clusters corresponded well to these experimentally identified structures, validating our single-chromosome modeling approach.

### SFIs compact the chromosome

To assess differences in chromosome compaction between the null and specific-fold models, we calculated pairwise spatial separations between chromosomal loci across various genomic distances in the specific-fold model (**Fig. 3A**, purple curve) and compared them with those from the null model (**Fig. 3A**, brown curve). We found that incorporating SFIs generally reduced spatial separations, especially at short genomic distances (<1000 Kb) (**Fig. 3A**), indicating a more compact overall chromosome structure. At larger genomic separations (∼ 2000 Kb), the specific-fold model showed slightly greater but not statistically significant mean distances than the null model (**Fig. S3A**). Importantly, the variability around the mean in spatial distances was much higher in the specific-fold model than in the null model, implying that SFIs disrupt the homogenous polymer behavior of the null model and increase chromosome-wide conformational heterogeneity.

**Figure 3:**
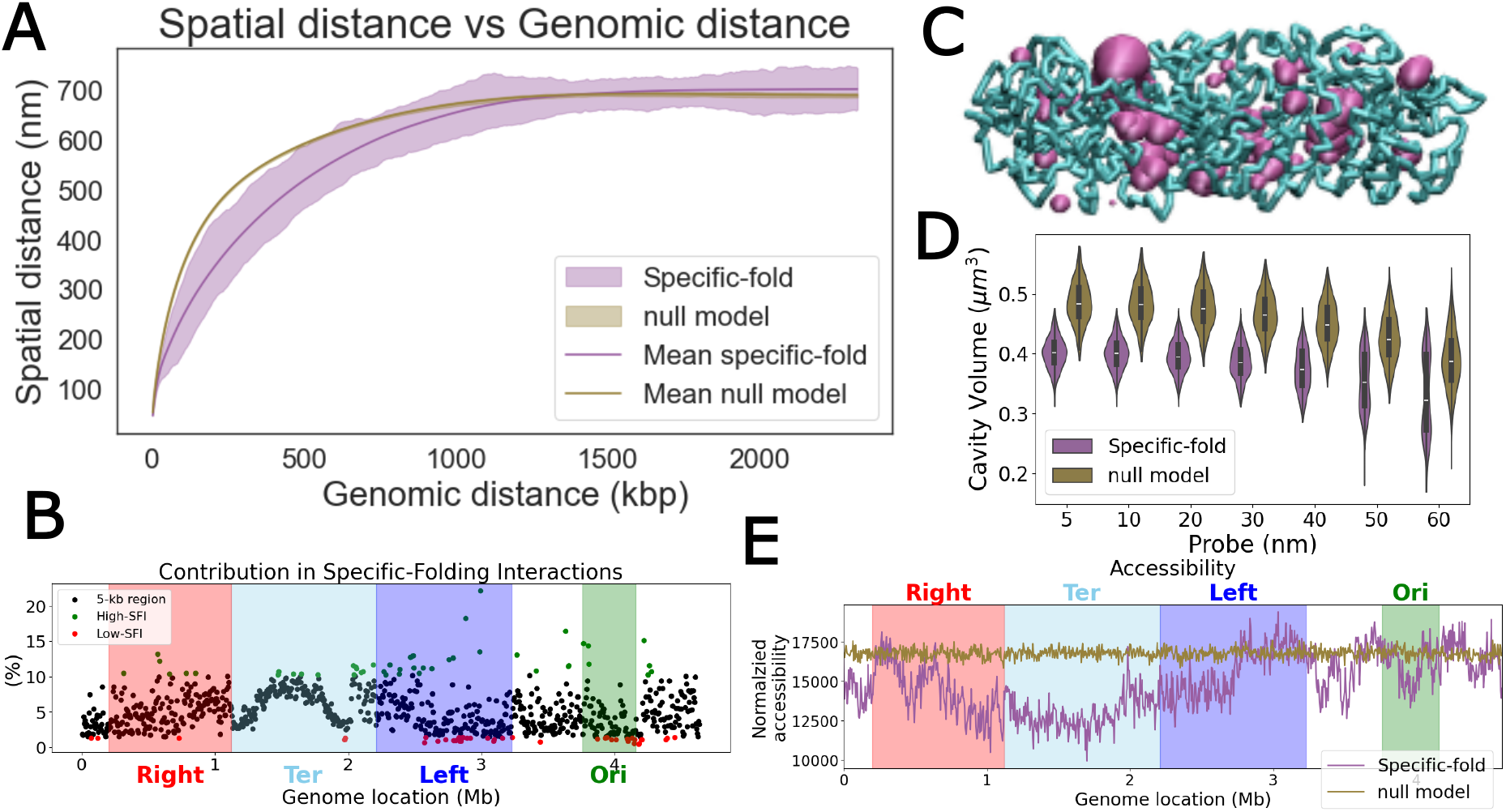
Effects of specific-folding interactions (SFIs) on chromosome topology and nucleoid accessibility. **A**) Mean spatial distance as a function of genomic distance showed that the specific-fold (purple) model had smaller spatial distances than those of the null model (brown) at short genomic distances. **B**) Percentages of SFIs in each 5 kb bin (black dots) along the chromosome. Bins containing the top 5% (green) or bottom 5% (red) SFIs were labeled. **C**) A representative example of a single, specifically folded chromosome (green) and the associated nucleoid-accessible regions (pink). **D**) Violin plots of the volumes of nucleoid-accessible regions (y axis) for probes of different sizes (x axis) for the specific fold model (purple) and the null model (brown). **E**) The specific-fold model (purple) showed variable nucleoid accessibility across different genomic regions compared to the null model (brown).

**Figure 4:**
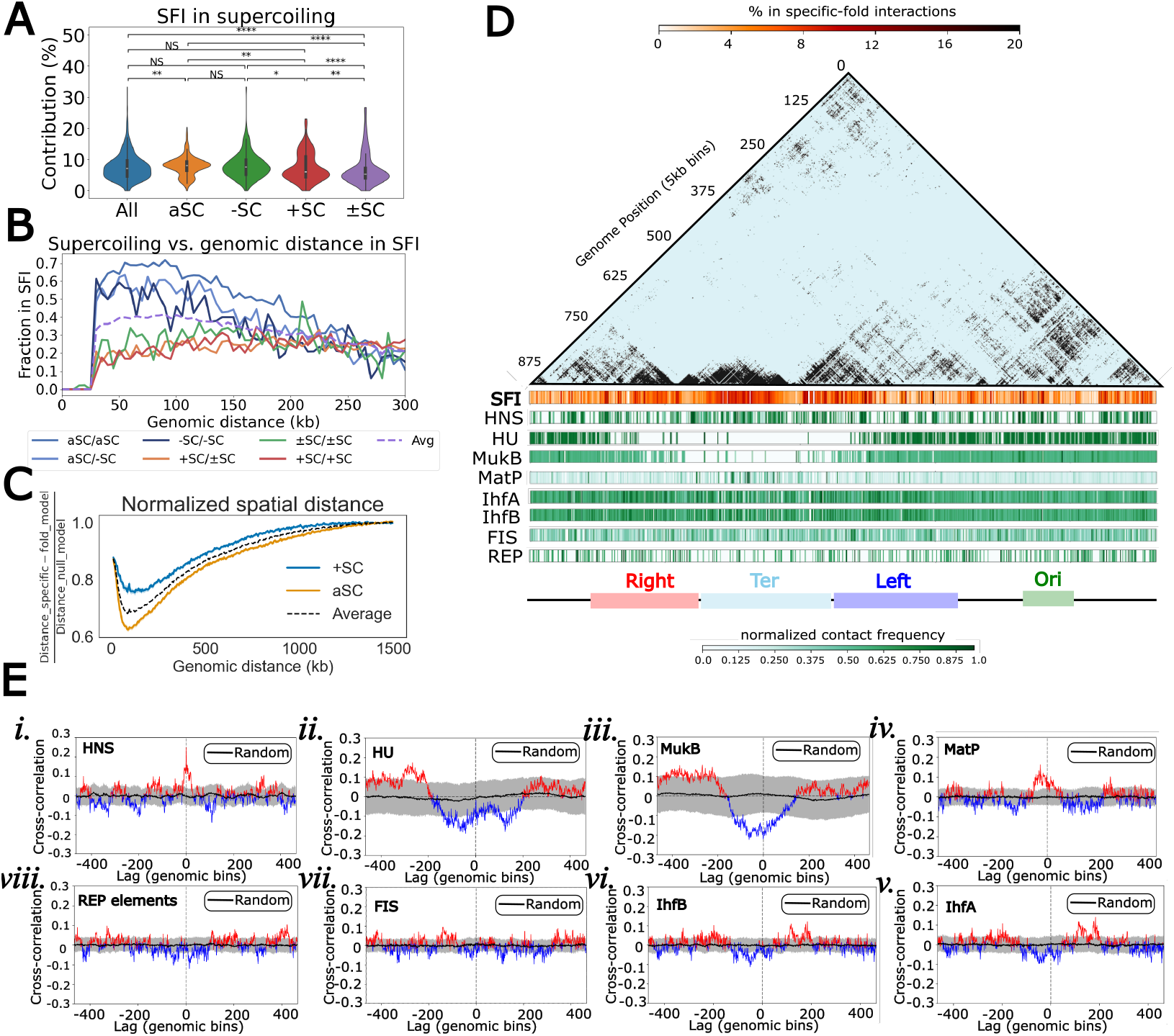
Contributions of supercoiling and nucleoid-associated proteins (NAPs) to specific-folding interactions. **A**) Violin plots of SFI percentages show that bins containing only +SC (positive supercoiling, red) had significantly lower SFIs than bins containing only -SC (negative supercoiling, green) and aSC (absence of supercoiling, orange), but higher SFI percentages than bin pairs containing both supercoils (±SC, purple). Statistical significance between groups is indicated as: **** (p < 1×10^−4^), *** (p < 1×10^−3^), ** (p < 1×10^−2^), * (p < 0.05), and NS (not significant). **B**) Percentages of SFI in chromosomal bin pairs of different supercoiling combinations across genomic separations. The dashed curve indicates the average SFI across all bin pairs, regardless of their supercoiling content. **C**) +SC-containing bin pairs showed relatively higher pairwise spatial distance than aSC bin pairs across all genomic separations. **D**) Distributions of SFIs and ChIP-seq data of various DNA-binding proteins and DNA elements in 5-kb bins across chromosome positions. The upper triangle shows SFIs as a binary map, with black indicating the presence of an SFI and light blue indicating its absence. The red colormap of SFI below the triangle indicates the total percentage of SFIs in that bin (see **Figure 3B**). The ChIP-seq maps below show the MACS2-calculated normalized enrichment of that protein within 5 kb bins (green colormap). For REPs, the map shows the number of REP elements within each bin. **E**) Cross-correlation functions between ChIP-seq protein binding values and SFI percentages at each bin. Values above zero (positively correlated) are colored in red, and values below zero (negatively correlated) are colored blue. The average cross-correlation functions of the null model with each protein’s ChIP-seq data are shown in black, with randomly permuted SFI and ChIP-seq peaks shown in gray as a control.

### SFIs are differentially distributed across the chromosome

To understand how SFIs were distributed along the chromosome, we calculated the percentage of SFIs for each chromosomal bin, defined as the proportion of SFIs observed in that bin divided by the total number of possible interactions of that bin with the remaining 927 genomic bins (**Fig. 3B**). We then identified the top 5% of bins with the highest SFI percentages as high SFI regions and the bottom 5% as low SFI regions (**Fig. 3B**, green and red dots, respectively). Notably, the *ter* region (**Fig. 3B**, cyan band) was enriched for high-SFIs and depleted of low-SFIs, whereas the *ori* region (**Fig. 3B**, green band) showed the opposite trend, being enriched for low-SFIs. In contrast, the right and left macrodomains (**Fig. 3B**, red and blue bands respectively) showed more even SFI distributions, indicating that their structural organization differs from that of the *ori* and *ter* regions.

### SFIs reduce nucleoid accessibility

Different classes of macromolecules play distinct functional roles in the cell, and their ability to access the nucleoid depends strongly on both their physical size and the structural organization of the chromosome. Quantifying nucleoid accessibility is therefore important for understanding how cellular processes such as transcription and translation are spatially regulated. The organizational deviation of the specific-fold model from the null model suggested that SFIs might modulate nucleoid accessibility to large biomolecules such as RNA polymerase and ribosomal subunits, which range in size from ∼10 to 30 nm (40).

To test this possibility, we used Alpha shape analysis (28), which captures the geometric surface and the internal porosity of three-dimensional (3D) chromosomal conformations more accurately than traditional convex-hull approaches (29). We observed that the specific-fold chromosome (see an example in **Fig. 3C**) consistently exhibited a reduced accessible volume relative to the null model across all probe sizes from ∼2 nm to 60 nm (**Fig. 3D**), suggesting that SFIs significantly reduce the physical space available to macromolecules within the nucleoid.

To determine whether the reduction in nucleoid accessibility was uniform across the genome, we averaged exposed surface areas across all chromosome conformations (**Fig. S3B, C**) and plotted accessibility along genomic locations. In contrast to the null model, in which the accessibility was relatively uniform (**Fig. 3E**, brown curve), the specific-fold model showed reduced accessibility in the *ter* region but higher accessibility in the *ori* region (**Fig. 3E**, purple curve). These observations suggested that the *ter* region is more compact than *ori*, consistent with previous high-resolution imaging studies (41).

### Cryptic prophages coincide with highly SFI regions

Given that chromosomal regions differ markedly in both SFI enrichment and nucleoid accessibility, we next asked whether these organizational differences corresponded to functionally distinct genomic features. One intriguing class of chromosomal loci is cryptic prophages, remnants of bacteriophage genomes that, although typically transcriptionally silent, can influence host physiology and stress responses (42). Because their repression is essential to prevent deleterious activation, their genomic placement may be subject to architectural constraints.

By comparing the identified high-SFI and low-SFI regions with 5 kb bins annotated to contain cryptic prophages, we found that among the eleven known cryptic prophages in *E. coli* (42), seven were located within high-SFI regions, including CPS-53, DLP12, CP4-44, and CP4-5, and none were found in low-SFI regions (**Fig. S4A**). Three of these, Qin/Kim, Rac, and *dif* sites, overlapped with high-SFIs and were positioned near the *ter* region, which exhibited increased chromosome compaction and reduced accessibility as shown in **Fig. 3E**. The enrichment of cryptic prophage in high-SFI regions and absence in low-SFI regions were significant (p < 0.008; see Methods, **Fig. S4B**). The co-localization of cryptic prophages with structurally compact, low-accessibility chromosomal domains suggested that genome folding may help maintain their transcriptional repression under non-inducing conditions. Such spatial sequestration could serve as a structural mechanism to prevent accidental prophage activation, thereby supporting bacterial survival in fluctuating environments.

### Negatively supercoiled or relaxed regions are enriched for SFIs, whereas positively supercoiled regions are depleted of SFIs

Given the non-uniform distribution of SFIs across the chromosome, we next asked what molecular factors might underlie their formation. One candidate is DNA supercoiling, a key element of transcription (43–46) and a major contributor to large-scale chromosomal organization (47). To examine the relationship between supercoiling and SFIs, we compared the genomic distribution of SFIs with ChIP-seq datasets for GapR-binding (30) and TopoI-binding (31), which preferentially mark positively (+SC) and negatively (-SC) supercoiled regions, respectively.

We found that bins containing positive supercoils (+SC) showed significantly lower percentages of SFIs than bins lacking supercoiling signals (aSC) or containing negative supercoils (-SC) (**Fig. 4A**; compare the red violin plots with the green and orange ones). This trend held consistently across chromosomal aSC/aSC, -SC/-SC, or aSC/-SC bin pairs separated by less than 150 kb (**Fig. 4B**, curves in blue shades); at 50 kb genomic separation, approximately 70% of interactions between aSC bins were SFIs, whereas fewer than 20% of interactions between +SC bins were SFIs (**Fig. 4B**, curves in red shades).

To further characterize the spatial behavior of +SC bins, we examined their pairwise spatial separations normalized to the null model. Across all genomic separations, +SC bin pairs were more spatially distant from one another than aSC bin pairs, with the largest relative difference occurring at ∼100 kb (**Fig. 4C**). We previously showed that +SC bin pairs exhibited higher chromosomal interaction frequencies than aSC bins, a finding also reproduced in the specific-fold model (**Fig. S4C**). We reasoned that the seemingly contradictory observation, that

+SC bins maintained higher interaction frequencies despite being spatially farther apart than aSC bin pairs at the same genomic separations (**Fig. S4D**), might reflect dynamic, frequent interactions between +SC bins driven by the high conformational mobility of protein-free +SC DNA, rather than a uniformly compact, static structure, as we previously proposed (48).This possibility is also consistent with the observed enrichment of SFIs in aSC and -SC bins and their depletion in +SC bins, suggesting that SFI formation was unlikely to be driven by DNA mechanics alone. Additional molecular factors, such as nucleoid-associated proteins (NAPs), may preferentially bind to aSC and -SC regions, mediating the specific contacts observed in these regions.

### Protein binding underlies SFIs

We reasoned that some of the factors contributing to SFIs are likely protein-based, especially NAPs and other DNA-binding proteins that organize the chromosome conformation. These proteins include HU (SRA: SRR353962) (49), H-NS (SRA: SRR37261926-29) (50), IHF (SRA: SRR353930 for IhfA and SRR353957 for IhfB) (49), and FIS (SRA: SRR15402973) (51), as well as Structural Maintenance of Chromosomes (SMC)-related and *ter*-organizing proteins such as MukB (SRA: SRR1946750-52) (52) and MatP (SRA: SRR1946729-31) (53). These proteins differ in DNA-binding affinity, sequence or motif preference, and functional roles in chromosome organization, DNA mechanics, and transcription regulation. For example, the histone-like HU protein preferentially binds bent or kinked DNA structures with high affinities and bridges long-range contacts (54, 55); H-NS recognizes AT-rich regions, restricts short-range interactions, and contributes to gene silencing (56); IHF binds defined DNA sequence motifs to differentially silence gene expression across growth phases (57); FIS bends AT-rich DNA and enhances transcription of select genes (58). MukBEF predominantly associates with the *ori* region to compact the chromosome along its length, whereas MatP binds specifically to *matS* sites in the *ter* region, thereby locally excluding MukBEF (52, 53). Because these proteins create defined, locus-specific chromosomal contacts, identifying their contribution to SFIs is essential for linking chromosome folding patterns to underlying molecular mechanisms.

We first plotted the percentage of SFIs in each chromosome bin as calculated in **Fig. 3B** along the chromosome (**Fig. 4D**, SFI). We then overlaid genome-wide binding peak values for DNA-binding proteins from the corresponding Chromatin Immunoprecipitation (ChIP)-seq datasets (**Fig. 4D**, proteins). To quantify their relationships, we computed cross-correlation functions between SFI and each protein’s ChIP peak values, which report whether and at what genomic distance SFI and protein enrichment are positively or negatively correlated (**Fig. 4E**, red and blue regions, respectively).

The most significant positive cross-correlation between SFI and protein enrichment was observed with H-NS, which showed a sharp, large positive correlation at a lag distance of 0 kbp and little correlation elsewhere (**Fig. 4Ei**), suggesting that H-NS co-localizes with SFIs across the entire chromosome. In contrast, SFIs were negatively correlated with HU and MukB binding across a broad range of genomic distances (**Fig. 4Eii, iii**, blue regions), consistent with the known depletion of both proteins from the *ter* region. SFIs showed a strong, positive, and statistically significant correlation with MatP binding, with the cross-correlation decaying rapidly within ∼100 kbp of lag distance (**Fig. 4Eiv**, red regions), indicating that MatP is a major local determinant of SFIs concentrated in the *ter* region. IHF-A and IHF-B binding sites, which are largely absent from *ori*, displayed positive correlations with SFIs but with a characteristic shift of ∼180 kbp relative to SFIs (**Fig. 4Ev, vi**), suggesting that IHF binding sites follow a similar but spatially offset chromosomal distribution pattern. FIS peaks did not show significant cross-correlations with SFIs beyond the permutation control (**Fig. 4Evii**), suggesting that FIS makes little direct contribution to SFIs. Similarly, Repeat Extragenic Palindrome (REP) elements, previously implicated in chromosome organization (50) and transcription regulation (27), showed no significant correlation with SFIs (314 unique bins, ∼1–2% of the genome, **Fig. 4Eviii**), indicating that REPs are not specifically associated with SFI formation.

Taken together, these analyses identified H-NS as a major chromosome-wide determinant of SFIs and MatP as a major *ter*-localized determinant of SFIs, both acting through direct co-localization with SFI-enriched regions. IHF-A and IHF-B contribute to SFIs through a phase-shifted chromosomal distribution pattern, while depletion of HU and MukB from SFI-enriched regions further shapes the broader architecture of specific chromosome folds. FIS and REPs did not appear to influence the formation of SFIs between genomic regions.

### Genes in low-SFI regions are expressed at higher levels than genes in high-SFI regions

Since the supercoiling state and several DNA-binding proteins both contributed to SFIs and are known transcription regulators, we next examined whether SFIs also correlated with transcriptional activity across the genome. Using RNA-seq data from *E. coli* cells in early exponential phase (SRA: SRR2932637) (33), we quantified gene expression levels in genomic regions with varying SFI densities. We then constructed a two-dimensional (2D) histogram in which each 5-kb bin was represented by its mean gene expression level and its SFI percentage and used 2D kernel density estimation (KDE) to visualize their joint distributions (**Fig. 5A**). The KDE revealed a clear enrichment of highly expressed genes in low-SFI regions, whereas genes associated with high-SFIs were preferentially shifted toward lower expression levels. Consistent with this correlation, categorical analysis showed that more genes belonged to the low-SFI/High-Expression group than to the high-SFI/high-Expression group (377 vs. 286), while transcriptionally repressed genes were enriched in high-SFI regions (371 vs. 272 Supplementary **Table S2, Fig. S5A**). A genome-wide correlation analysis further identified a weak but statistically significant inverse relationship between gene expression and SFI density (*ρ*_*Spearman*_ = −0.088, *p* = 3.1 × 10^−10^), indicating that regions with more SFIs tend to be less transcriptionally active overall.

**Figure 5:**
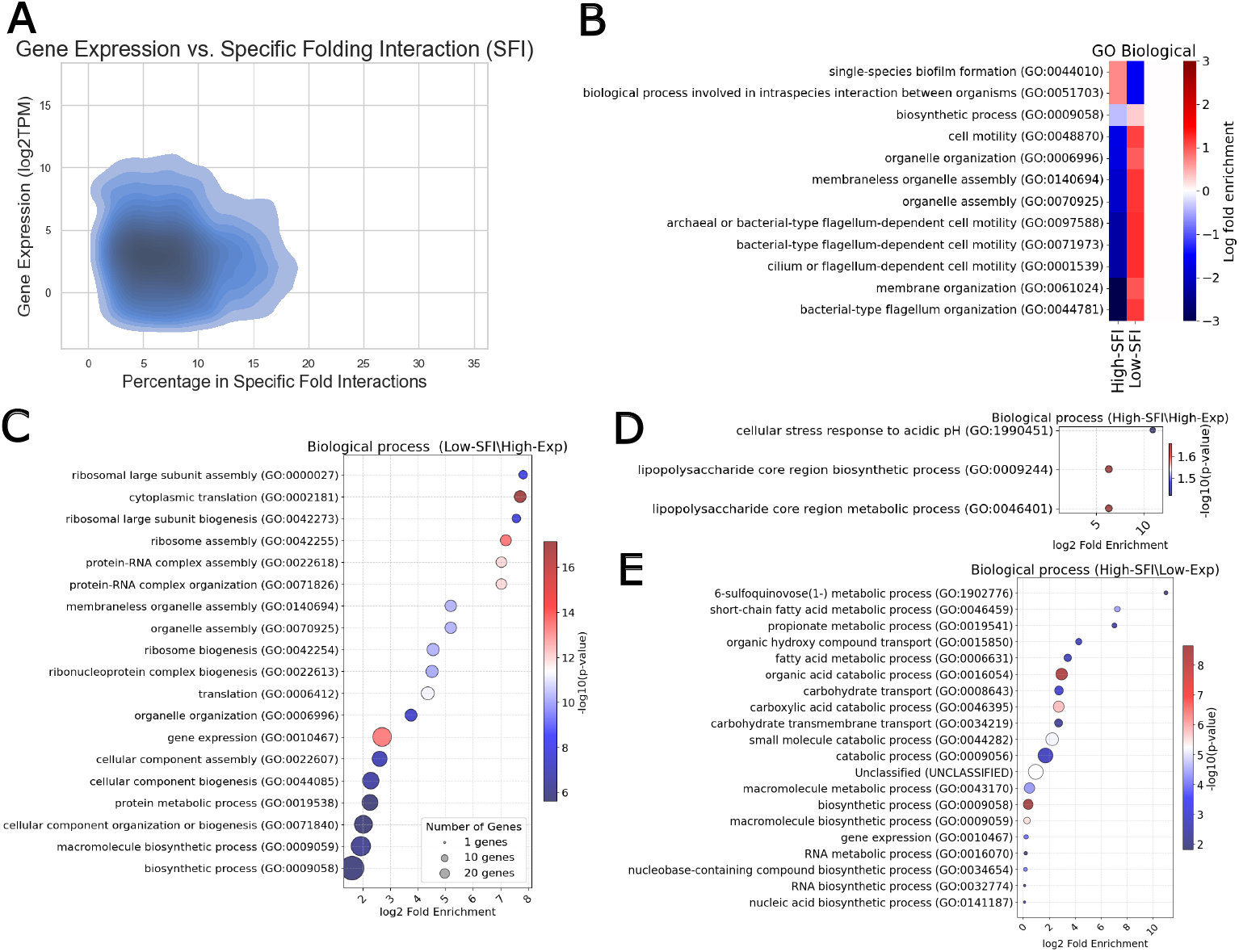
Gene ontology analysis reveals a functional partitioning of the chromosome where low-SFI regions predominantly harbor highly expressed genes involved in core processes, whereas high-SFI regions are biased toward lowly expressed genes involved in stress-response programs. **A**) 2D kernel density estimation for gene expression and the corresponding SFIs. **B**) Gene Ontology enrichment analysis using PANTHER for genes in high-SFI and low-SFI. **C**) Biological process of gene ontology of highly expressed genes in low-SFI regions. **D**) Biological process of gene ontology of highly expressed genes in high-SFI region. **E**) Biological process of gene ontology of lowly expressed genes in high-SFI regions.

We next compared mean gene expression levels across 5 kb bins classified as high-SFI (top 25%) and low-SFI (bottom 25%). A Mann-Whitney U test revealed that the average gene expression per bin in low-SFI regions was significantly higher than in high-SFI regions (p<0.001, **Fig. S5B)**. Consistently, when considering all individual genes within each bin, individual genes located in high-SFI regions had significantly lower expression than genes in low-SFI regions, also confirmed by a Mann-Whitney U test (*p* < 10^−8^, **Fig. S5C)**. These findings indicated that high-SFI regions were more strongly associated with reduced transcriptional activity, consistent with increased chromosomal compaction and decreased nucleoid accessibility, which may limit engagement of the transcription machinery.

### Gene ontology analysis reveals distinct patterns of gene expression linked to SFIs

Because SFI density correlated with gene expression levels, we next asked whether genes in different SFI regions also differed systematically in their biological functions. We performed a gene ontology (GO) enrichment analysis (59) to assess whether specific biological processes were statistically overrepresented in genes from high- or low-SFI regions.

We first compared all genes in high-SFI regions with those in low-SFI regions (**Fig. 5B**). In this whole-genome comparison, we found that high-SFI regions were enriched for GO terms associated with sessile and community-associated lifestyles, such as single-species biofilm formation (GO:0044010) and multi-organism processes (GO:0051703). In contrast, low-SFI regions were biased toward functions supporting a more motile, exploratory state, including cellular motility (GO:0048870, GO:0071973), flagellum assembly, membrane and organelle organization, and diverse biosynthetic processes (GO:0009058), consistent with a metabolically active, dispersive lifestyle.

We next examined GO enrichment specifically among highly expressed genes in low-SFI regions versus high-SFI regions. Highly expressed genes in low-SFI regions were enriched for broad, essential cellular functions, including macromolecule biosynthetic processes (GO:0009059), protein metabolism (GO:0019538), gene expression (GO:0010467), and ribosome biogenesis (GO:0042254) (**Fig. 5C**). These categories pointed to core housekeeping and growth-related functions that support homeostasis under steady-state conditions. In contrast, highly expressed genes in high-SFI regions were enriched for more specialized and stress-associated functions, such as cellular response to acidic pH and lipopolysaccharide biosynthetic processes, consistent with context-dependent regulation in challenging environments (**Fig. 5D**).

We also analyzed lowly expressed genes in high-SFI regions (**Fig. 5E**). These genes were enriched in fatty acid metabolic process (GO:0006631), short-chain fatty acid metabolic process (GO:0046459), propionate metabolic process (GO:0019541), organic acid catabolic process (GO:0016054), and carboxylic acid catabolic process (GO:0046395). The reduced expression of these pathways suggested decreased catabolic activity and limited utilization of carbon substrates, consistent with a more compact, transcriptionally repressed chromatin state in high-SFI regions.

Taken together, these results supported a functional partitioning of the chromosome: low-SFI regions, which are more accessible and less compact, predominantly harbor highly expressed genes involved in core biosynthetic and regulatory processes, whereas high-SFI regions, which are more compact and less accessible, are biased toward genes mediating sessile, stress-resistant, and community-associated phenotypes. This division aligns with the enrichment of cryptic prophages in high-SFI domains and supports a model in which chromosome folding and accessibility help segregate core cellular functions from conditionally activated stress-response programs.

## Conclusion

A central challenge in understanding bacterial chromosome organization is distinguishing biologically encoded specific-folding interactions (SFIs) from the nonspecific compaction that inevitably arises from confining a circular polymer within the nucleoid. In this study, we developed a polymer-based simulation framework for the circular *E. coli* chromosome in a confined, capsule-shaped nucleoid and showed that, by separating polymer-driven, nonspecific contacts from biological SFIs, we recapitulated the global features of experimental Hi-C maps while uncovering structural features not explained by the polymer’s confinement alone. The specific-fold model reproduced chromosomal interaction domains (CIDs), captured insulation boundaries, and revealed substantial heterogeneity among single-cell conformations, consistent with experimental imaging results. SFIs reshaped chromosome architecture by increasing compaction, reducing nucleoid accessibility, and differentially organizing large-scale regions. In particular, the *ter* region was more compact and enriched for high-SFI interactions, whereas the *ori* region remained more accessible. High-SFI regions were also strongly associated with cryptic prophages, linking specific folding to stress-related genomic elements.

Our results showed that the supercoiling state is closely associated with SFIs. Bins containing positive supercoils (+SC) exhibited significantly lower SFI enrichment, whereas bins lacking supercoiling (aSC) or containing negative supercoils (−SC) were markedly enriched in SFIs. At finer scales (∼50 kb), interactions among aSC bin pairs were largely dominated by SFIs, whereas +SC bin pairs contributed only a small fraction of SFIs. This depletion of SFIs in +SC regions persisted across genomic separations and coincided with increased spatial separations. These results suggested that +SC regions form frequent, dynamic interactions rather than participating in cohesive long-range folding, whereas aSC and −SC regions preferentially engage in SFI-driven interactions.

Furthermore, our results indicated that SFIs reshaped chromosome topology and produced heterogeneous genome-wide accessibility, with particularly pronounced effects in the *ori* and *ter* macrodomains. The heterogeneous accessibility patterns observed in our model are consistent with structural heterogeneity previously reported for the *E. coli* chromosome, where nucleoid-associated proteins (NAPs) organize DNA into macrodomains and locally compacted regions (4). Sequence-dependent binding of these proteins, together with genetic features distributed along the chromosome, may therefore impose spatially heterogeneous structural constraints that favor the formation of specific contacts detected in Hi-C. In addition, the bacterial chromosome behaves as an active polymer, in which ATP-driven processes, including loop extrusion by Structural Maintenance of Chromosomes (SMC) complexes and transcription-associated forces generated by RNA polymerase, can dynamically reorganize its structure. Such active processes may selectively reinforce or stabilize subsets of SFIs, thereby further contributing to the differential accessibility patterns captured by our model.

Finally, gene ontology analysis highlighted functional differences between high-SFI and low-SFI regions. Low-SFI regions were enriched for core cellular and biosynthetic functions and exhibited higher transcriptional activity, whereas high-SFI regions were associated with more specialized processes linked to environmental response and stress adaptation. These findings align with the strong overlap between high-SFI regions and cryptic prophage loci, supporting the idea that high-SFI regions may contribute to community-associated, stress-resistant phenotypes.

Together, our findings show that although polymer physics accounts for much of *E. coli* chromosome folding, SFIs superimposed on this background play a central role in shaping domain structure, accessibility, and functional genome organization. More broadly, this framework provides a quantitative basis for dissecting how biologically encoded interactions contribute to three-dimensional chromosome organization. It can be extended to other bacterial species and used to test causal effects through *in silico* perturbations.

## Acknowledgements

*Author Contributions*: P.D. conceived the study with J. Liang and J. Xiao; P.D. designed and performed the majority of the computational modeling, software development, data analysis, validation, and visualization. T.N. contributed to formal analysis and investigation of protein binding data, visualization, and manuscript editing. H.F. assisted with formal analysis, visualization, and interpretation of results. F.H., L.D., and J. Liu contributed to manuscript editing and review. A.M. and J.X. provided resources, project supervision, administrative support, critical feedback throughout the study, and manuscript writing. J. Liang led the conceptual framework, secured funding, and supervised the overall project. P.D. wrote the original draft, with all authors contributing to the review and editing of the manuscript.

## Code availability

C++ source code for circular chromatin polymer folding, along with tutorials for configuring and running polymer folding simulations, will be made available upon reviewer request and will be publicly released before the final revision.

