## Supplementary Information for "Disentangling polymer confinement from specific-folding interactions reveals the drivers of *E. coli* chromosome organization"

* Corresponding authors

#### Supplementary figures

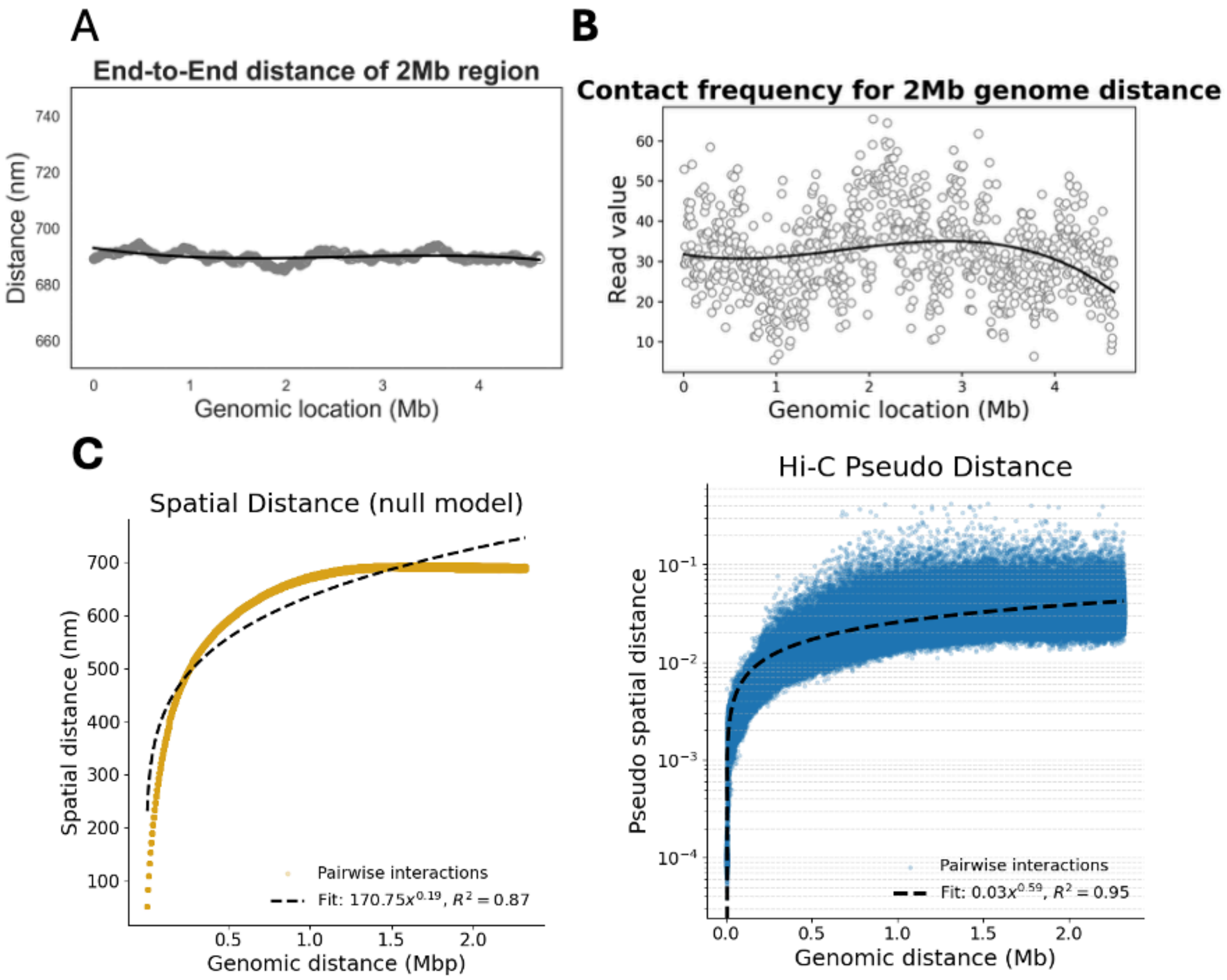


**Figure S1. null model explains the majority of global patterns observed in Hi-C. A)** The average end-to-end distances of 2 Mb chromosomal segments across various genomic locations in the null model remain nearly identical, indicating unbiased chain conformations regardless of bead position. **B**) The contact frequency between chromosomal loci separated by 2 Mb in the experimental Hi-C data shows a nearly uniform mean across the genome, although certain regions exhibit notable deviations from this average due to local chromatin interactions. **C**) The mean spatial distance of the null model (left) and the pseudo spatial distance (reverse of contact frequency) of the experimental Hi-C data (right) as a function of genomic separation show that the spatial distance scales as a power-law of genomic distance (exponents = 0.19 and 0.59, respectively). Both relationships exhibit power-law scaling, although the Hi-C-derived pseudo distance should not be directly interpreted as a Euclidean spatial distance, which likely contributes to the difference in scaling exponents.

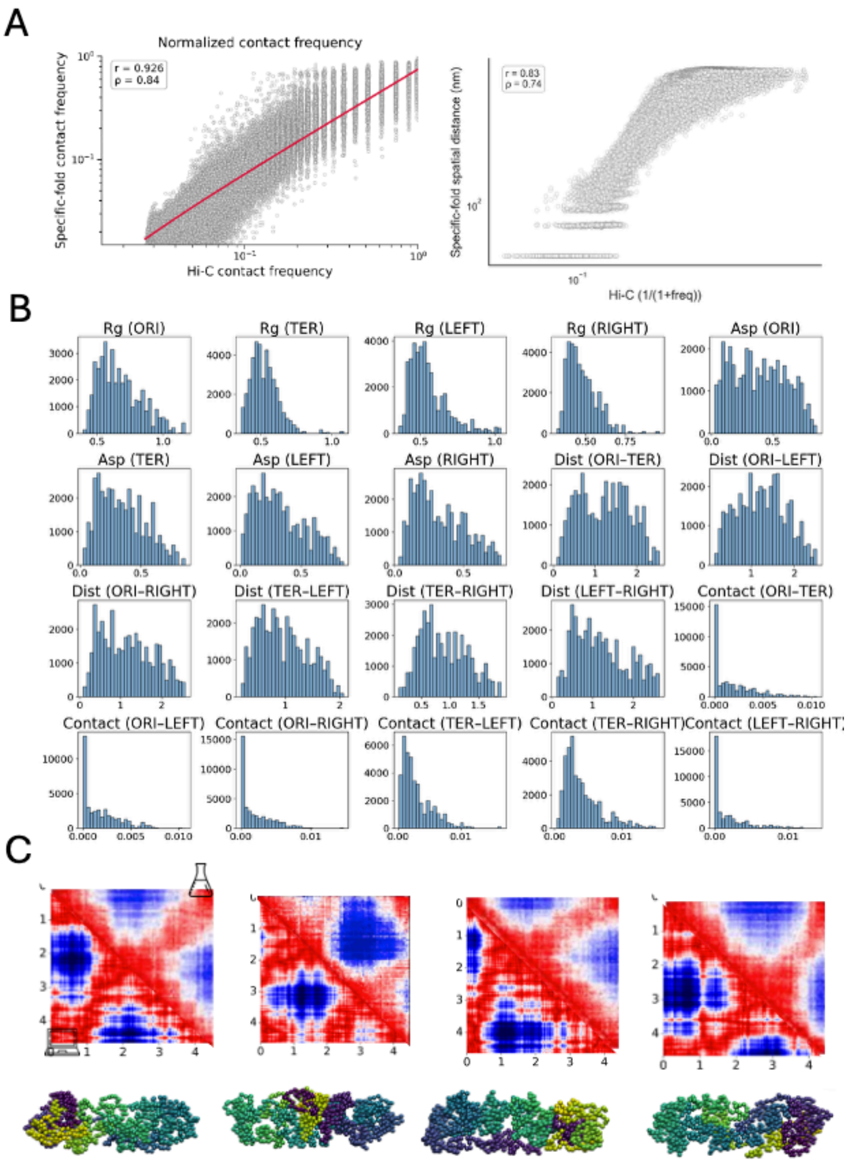


**Figure S2. Single-cell specific-fold model recapitulates specific local interactions. A**) Contact frequencies between experimental Hi-C and specific-fold models (left panel) and the average spatial distance calculated from the null model and the Hi-C–derived distance (right panel) are highly correlated. The Spearman correlation coefficients are 0.83 and 0.74, respectively. **B**) Geometric properties of specific fold model across single-cell conformations. Histograms of the radius of gyration for the *ori*, *ter*, *left*, and *right* chromosomal regions are shown. Asphericity distributions indicate that the *ter*, *left*, and *right* regions adopt more spherical conformations compared with the *ori* region. Pairwise distances and contact frequencies among the *ori*, *left*, *ter*, and right regions are quantified, showing that the *ter* region interacts more frequently with the *left* and *right* regions than any other pairwise combination. C) Pairwise distance maps from simulations and experiments show consistent folding patterns. The upper triangle represents simulation-derived pairwise distances, and the lower triangle represents experimental measurements (1). Representative 3D structures corresponding to each conformation are shown below the heatmap.

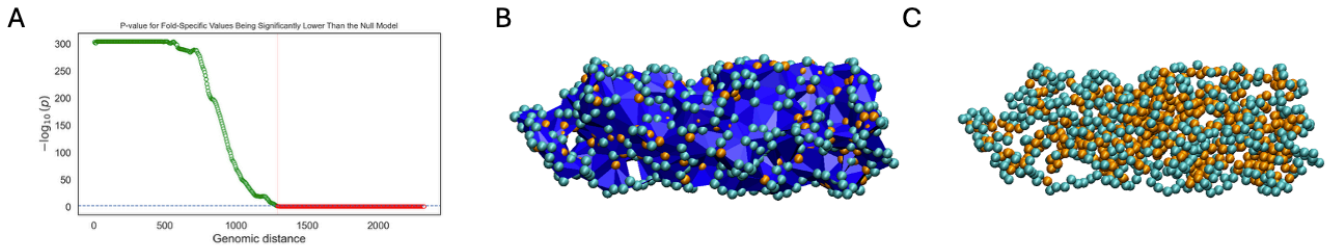


**Figure S3**. **Geometrical analysis of the specific-fold and null models. A)** One-sided Mann–Whitney p-value test to assess whether the spatial distances between two loci of different genomic distances in the specific fold model are closer than those of the null model. At ~1200 kb separation (dashed line), the differences between those of the specific fold model are no longer significantly closer than those of the null model. **B**) Wrap (blue) calculated by alpha shape for the chromosome, with accessible regions of the chromosome shown as cyan beads. **C**) Accessible region of the chromosome shown as cyan beads. Orange beads correspond to inaccessible surfaces in the chromosome.

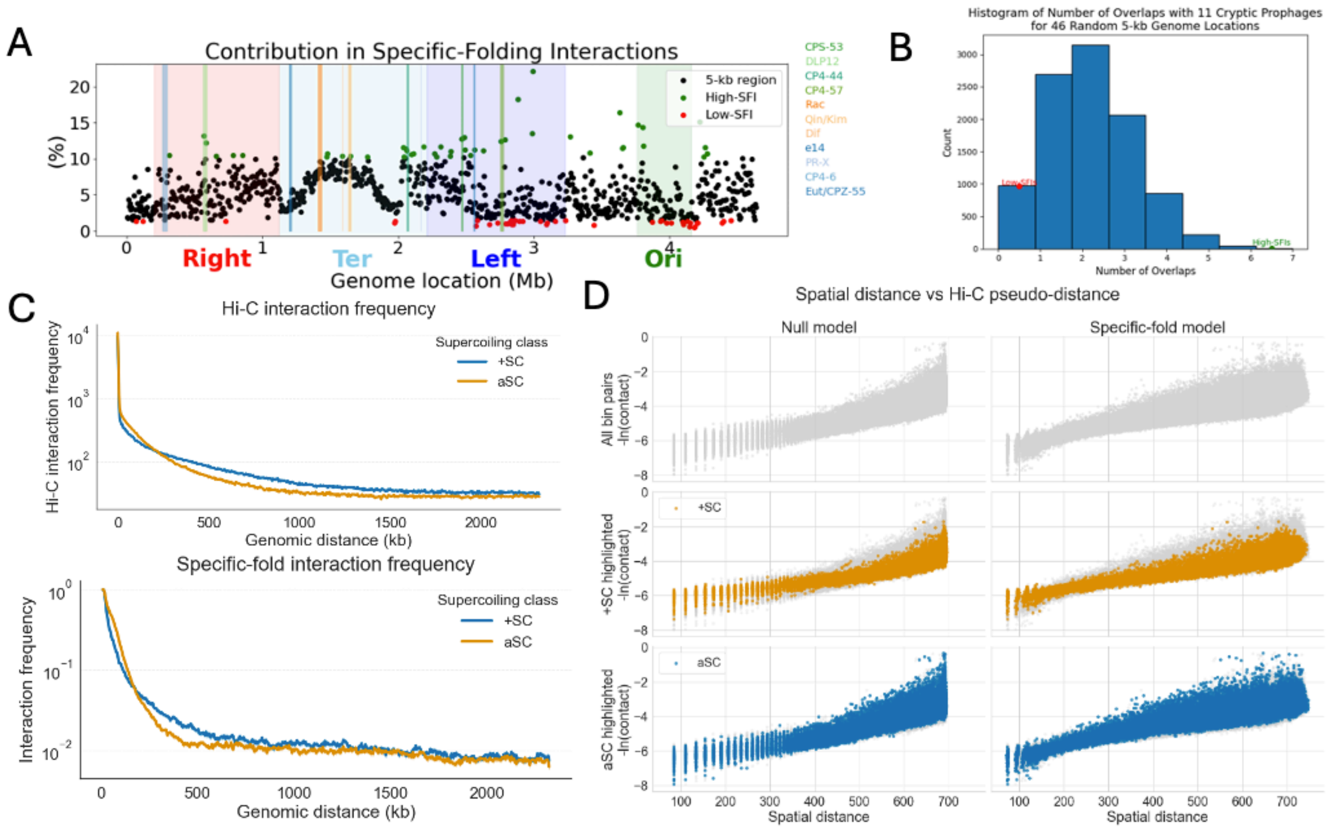


**Figure S4**. **SFI is associated with cryptic prophages, supercoiling, and protein-binding sites. A**) Genomic locations of cryptic prophages (vertical lines). Top (green) and bottom (red) 5% SFIs 5-kb genomic bins (black) are indicated. **B**) The enrichment of cryptic prophages in high SFIs and absence in low SFIs is significant. Permutation-based null distribution of overlaps with 11 cryptic prophage regions, generated by randomly selecting 46 regions (5% of 928 bins) across 10,000 iterations. The histogram shows the expected number of overlaps with the 11 regions. The red dots indicate the observed overlap for the low-SFI regions, while the green dot indicates the observed overlap for the high-SFI regions. **C)** Interaction frequency averages for +SC and aSC bin pairs in the Hi-C experiment (top) and the specific-fold model (bottom). Both show similar interaction frequency patterns, with +SC bin pairs exhibiting higher interaction frequencies at genomic separations greater than ~250 kb. **D)** Pairwise spatial distance and pseudo Hi-C distance (−ln (contact frequency)) (top panels) with overlays highlighting +SC (middle panels) and aSC bin pairs (lower panels). +SC bin pairs exhibit lower pseudo Hi-C (higher interaction frequencies) at similar spatial distances, particularly around ~400 nm, suggesting enhanced detectability of +SC interactions in Hi-C experiments.

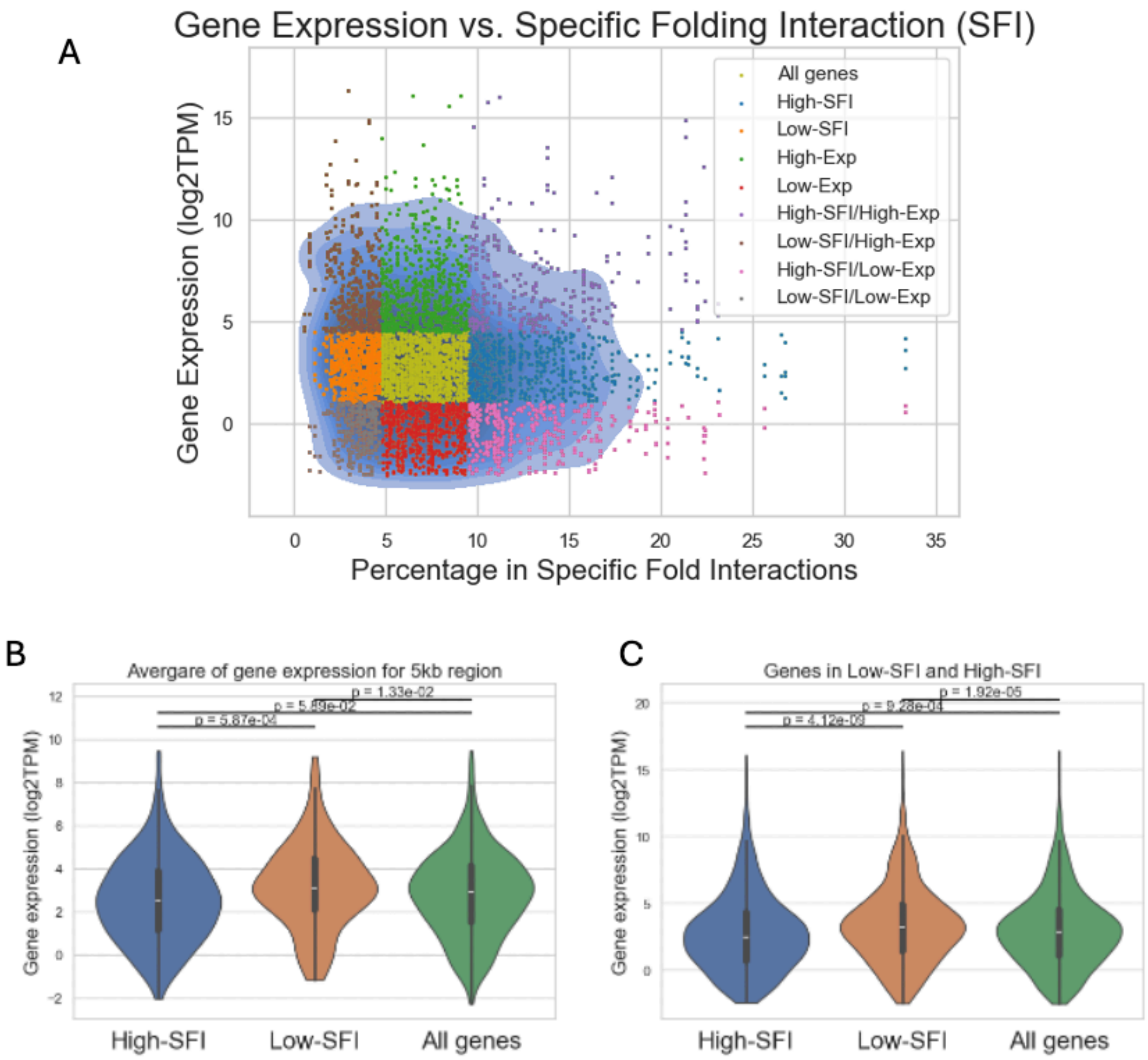


**Figure S5. SFI is associated with low gene expression. A)** Genes were categorized into three comparison groups: 1) High-SFI vs. Low-SFI, 2) High-SFI/High-Exp vs. Low-SFI/ High-Exp, and 3) High-SFI/Low-Exp vs. Low-SFI / Low- Exp. **B)** Average gene expression of regions with the highest and lowest densities of SFI (Top 25 are High-SFI and Bot 25 are Low-SFI) **C)** Gene expression of gene in regions with least and most SFI

### Supplementary tables

**Table S1:** Cluster membership of single-cell conformations

| Cluster No | 1 | 2 | 3 | 4 | 5 | 6 | 7 | 8 | 9 | 10 | 11 | 12 |
|---|---|---|---|---|---|---|---|---|---|---|---|---|
| Population (%) | 7.25 | 5.25 | 8.75 | 8.25 | 7.80 | 12.00 | 9.60 | 10.15 | 6.25 | 7.95 | 10.25 | 6.50 |

**Table S2:** Number of genes in each quadrant

| # of Genes | | SFI | | | |
|---|---|---|---|---|---|
| | | Low | Mid | High | Total |
| Expression | Low | 272 | 636 | 371 | 1,279 |
| | Mid | 628 | 1323 | 608 | 2,559 |
| | High | 377 | 616 | 286 | 1,279 |
| | Total | 1,277 | 2,575 | 1,265 | 5,117 |
